# Optimal data-driven parameterization of coiled coils

**DOI:** 10.1101/353532

**Authors:** Dmytro Guzenko, Sergei V. Strelkov

## Abstract

α-helical coiled coils (CCs) represent an important, highly regular protein folding motif. To date, many thousands of CC structures have been determined experimentally. Their geometry is usually modelled by theoretical equations introduced by F. Crick that involve a predefined set of parameters. Here we have addressed the problem of efficient CC parameterization from scratch by performing a statistical evaluation of all available CC structures. The procedure is based on the principal component analysis and yields a minimal set of independent parameters that provide for the reconstruction of the complete CC structure at a required precision. The approach is successfully validated on a set of canonical parallel CC dimers. Its applications include all cases where an efficient sampling of the CC geometry is important, such as for solving the phase problem in crystallography.

## 1. Introduction

α-helical coiled coils (CCs) representing a superhelical arrangement of two or more α-helices are an important protein structure motif [1]. As much as 8% of all human genome proteins are estimated to include CC motifs [2]. The basis of the CC formation is the favourable interaction of α-helices that show a periodic pattern (most typically, a heptad) in the hydrophobicity of the residues along their sequence [3]. Early on, an analytical representation of a CC structure through a set of simple equations was proposed [4]. This representation assumed a regular and symmetric arrangement of the α-helices forming the CC. Its parameters included the radius and pitch of the superhelix, the phase of the first residue (often expressed by the so-called Crick angle, also known as the interface angle [5]), and the radius, pitch and rise per residue of the α-helix [6]. In the past, two main approaches to derive the set of Crick parameters from a given experimental structure were proposed. The first approach was based on a best fit between a theoretical structure parametrized through Crick equations and the experimentally determined CC [7], [8]. The second approach involved a direct measurement of the local Crick parameters along the structure using the program Twister [6]. While the Crick parameterization is a mathematical abstraction, for naturally occurring CCs the corresponding parameter space is quite restricted with some parameters clearly correlated [8]. At the same time, additional parameters beyond the Crick representation, such as the axial offset of individual α-helices forming the CC for example, are sometimes introduced to better describe the geometry of natural CCs [9].

In this manuscript, we reconsider the problem of optimal CC parameterization from scratch *i.e.*, not assuming any predefined analytical representation. Our goal is to obtain the most accurate description that would use a minimal number of parameters. To this end, we analyze a large set of experimentally determined CC structures and infer the relevant structural parameters in a statistically rigorous way. This is done through the principal component (PC) analysis of all such structures, whereby the orthogonal PCs are derived without the need to explicitly define them. The given CC structure is then described by a set of PC scores corresponding to the amplitudes of deviations from the mean CC structure. Since the PCs can be ranked by their ’importance’ (i.e., the explained variance ratio in the dataset), a certain number of top-scoring parameters can be included towards achieving the required modelling accuracy with respect to the experimental structure. While we will focus the description of our algorithm on parallel CC dimers, other CC structures are treated in the same principle way. The present code implementation can handle parallel and antiparallel dimeric CC structures, as well as trimers.

## 2. Sets of experimental α -helical and CC fragments

As a starting point, all parallel dimeric CC fragments of 15 residues per chain were extracted from the CC+ database [10] and truncated to polyalanine. Following the procedures described in [11], a non-redundant set of 14175 CC fragments was obtained. Here non-redundancy was defined in terms of stuctural dissimilarity, so that the root mean square deviation (RMSD) between any two fragments upon their optimal superposition was at least 0.2Å. A non-redundant set of 15760 15-residue α-helices with a minimal pairwise RMSD of 0.1 Å was constructed in a similar way. All RMSD values for 15-residue α -helices and CC fragments given here and below are for all-atom polyalanine models.

## 3. PC analysis and parameterization of an α-helix

Multiple structural alignment of the non-redundant set of 15-residue α-helices was obtained using iterative superposition method [12]. Thereafter the aligned coordinates were subjected to a PC analysis. This resulted in a decomposition of each α-helix into a linear combination of the PC vectors:

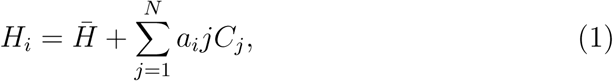

where 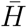 is the mean α-helix, *C*_*j*_ is the *j*-th component and *a*_*j*_ is its score.

Each PC represents a mode of variation of the α-helical geometry, with the explained variance in the dataset proportional to the eigenvalues of the coordinate covariance matrix (Fig. 1a). The resulting three dominant modes including two perpendicular ’bend’ modes (Fig. 1b) and one ’twist’ (Fig. 1c) mode and their variance ratios are consistent with those previously reported [13]. The encoding system consisting of the mean helix 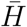 and scores of the first three components allows to model individual α-helices from the non redundant set with a mean RMSD of 0.23Å (Fig. 1d).

**Figure 1:**
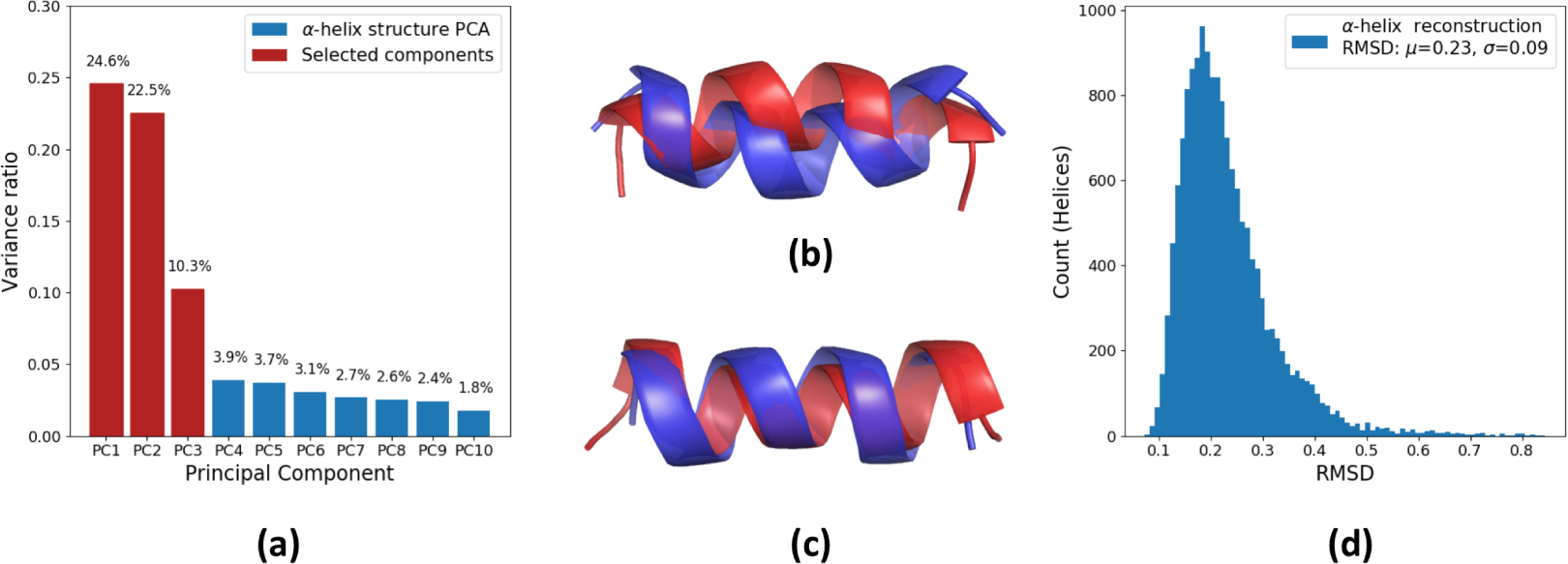
Principal component analysis of a 15-residue α-helix. (a) Variance ratio per PC of all α-helices in the non-redundant set. (b) The first PC corresponds to the bending of the α-helix in one direction. The second PC represents bending in a perpendicular direction and is not shown. The third PC corresponds to over/underwinding of the α-helix (exaggerated in the figure). (d) Distribution of all-atom RMSD values calculated by superposing model α-helices encoded by the three PC parameters with the original α-helices.

## 4. Extended parameterization and initial analysis of the CC structure space

To approach parameterization of a CC structure (of any multiplicity), we first align each of its α-helices to the mean α-helical structure 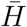. The result of such alignment for each helix is reflected in the scores of the three dominant PCs described above (Fig. 1a). Next, we record the position of the second, third *etc*. helix with respect to the first one. This position is given by the rotation matrix *R* and the translation vector *T* (Fig. 2a). The description of a CC structure is then given by:

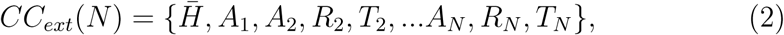

where *N* is the number of helices in the CC, *A*_*i*_ is the vector of PC scores for the *i*-th α-helix, *R*_*i*_ and *T*_*i*_ are rotation and translation parameters of the *i*-th α-helix.

**Figure 2:**
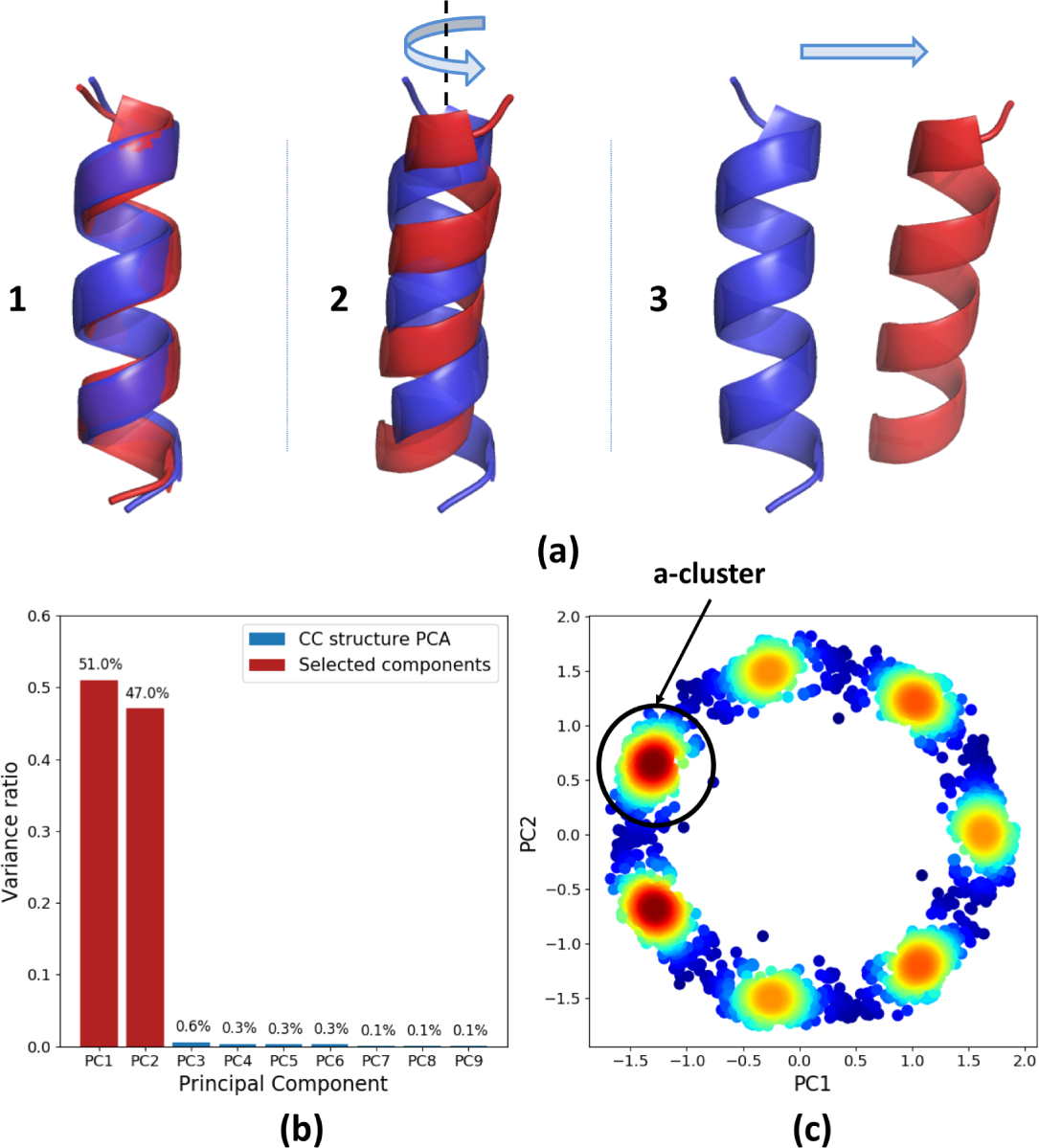
Initial clusterization of dimeric CC fragments. (a) CC extended parameterization procedure. (1) Start with an average helix and apply α-helical PC parameters. (2) Rotate and (3) position the second α-helix relative to the first one. (b) Variance ratio per PC of all CCs in the non-redundant set. (c) The CCs projected into the space of their first two PCs. Colour indicates the density of the points from low (blue) to high (red).

For a CC dimer this process results in an ’extended set’ of 18 parameters including three parameters for each of the two α-helices, nine parameters for the rotation matrix and three for the translation vector of the second helix.

Next, we perform a PC analysis on the non-redundant set of 15-residue dimeric CC fragments, each represented by a vector of extended parameters from eq. 2. The resulting ranking indicates a clear dominance of the first two PCs (Fig. 2b), whereby seven pronounced structural clusters in the space of these two PCs are observed (Fig. 2c). This pattern reflects the fact that the fragment set is dominated by the structures with regular heptad repeats which can start at one of the seven possible heptad positions. The two most prominent clusters correspond to the fragments starting with either *a* or *d* heptad position. This is easily explained by the fact that all fragments recorded in the CC+ database start and end with a core residue.

## 5. Optimal parameterization of 15-residue CC dimers

As the next step, we focus on a cluster of 15-residue fragments that all have the same heptad register, *e.g.*, those starting at position *a.* Towards this end, all fragments situated in the vicinity of one of the maxima in the space of the first two PCs (2c) are selected. Since the initial non-redundant set included overlapping 15-residue fragments, this cluster of fragments alone is structurally representative of all sufficiently long dimeric CCs in the CC+ database.

Results of the PC analysis of this cluster are presented in Fig. 3. The average structure of the cluster is expectedly a canonical left-handed symmetric CC. To model a CC structure with a desired number (*k*) of parameters, the first *k* PC vectors are used, and the respective variations are added to the mean parameters vector (eq. 2). Technically, the rotation matrix of an individual α-helix restored with this procedure, 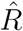, may no longer be orthogonal. Therefore we use the nearest orthogonal matrix as in [14] instead:

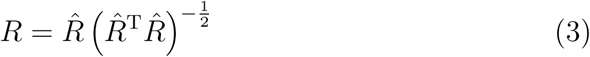

**Figure 3:**
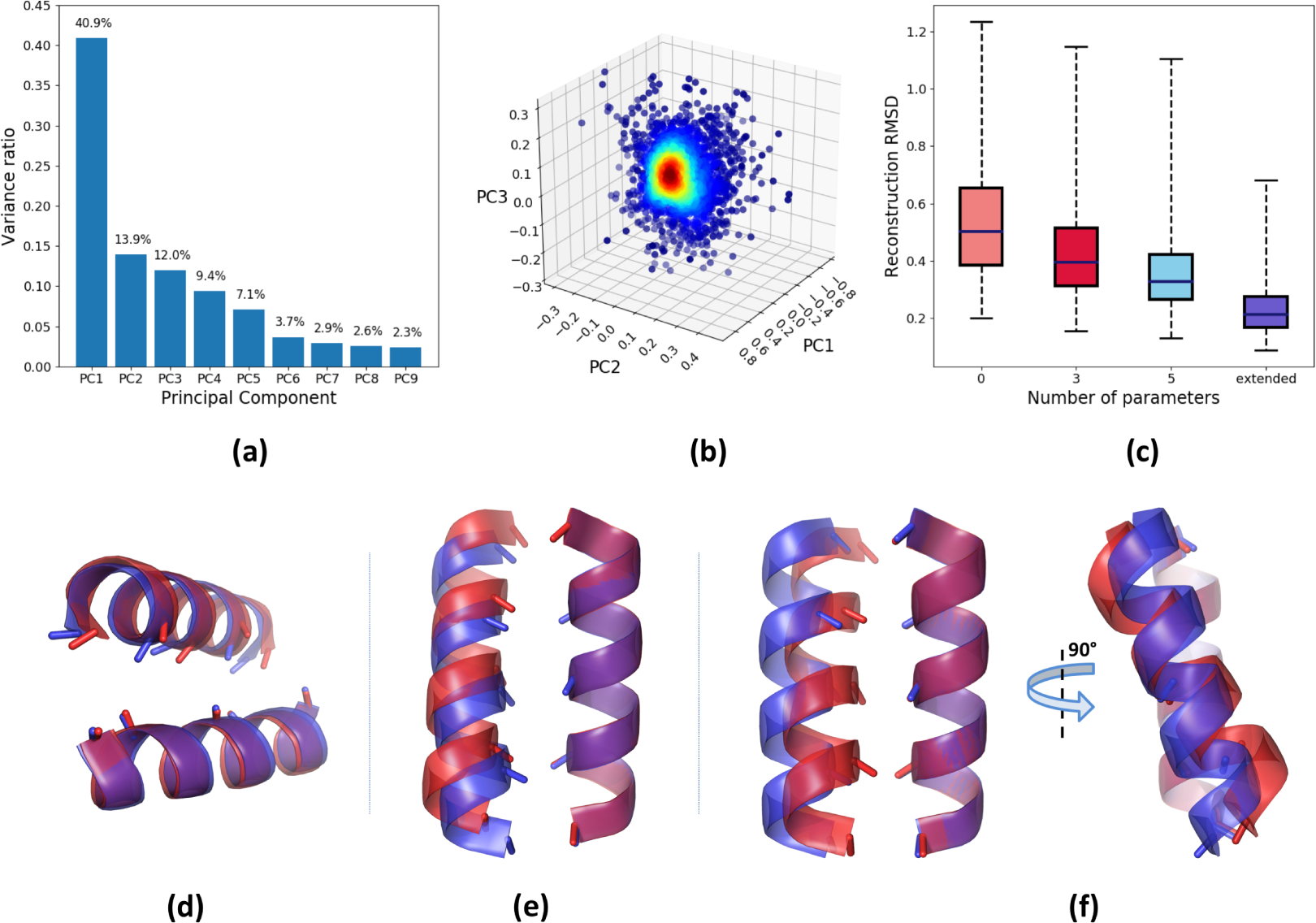
Dimensionality reduction of the a-cluster of CC fragments. (a) Variance ratio per PC of all selected CC fragments. (b) The CCs projected into the space of their first three PCs. Colour indicates density of the points from low (blue) to high (red). (c) Box plots of the RMSD distributions for the CC fragments aligned to a mean structure (light red), represented with the 3 (dark red) and 5 (light blue) PC parameters and with the extended parameterization (dark blue). Each box plot shows minumum and maximum values (bars), middle 50% of the data (box) and the median value (line). (d,e,f) Variations of the CC geometry corresponding to the first (d), second (e) and third (f) obtained PCs. Structures resulting from negative (blue) and positive (red) score values are superposed for each case.

Next, we wanted to explore how efficiently the space of all 15-residue dimeric CC structures can be sampled using a particular number of parameters. To this end, for each structure in the chosen cluster of experimental CCs, we have built the best-matching model upon varying the first *k* PC parameters. The resulting statistics is presented in Fig. 3c. A zero-PC modelling (which corresponds to simply approximating any structure with the cluster average) results in a median RMSD of 0.5Å, with a maximal RMSD value observed being 1.2Å. These relatively low values (obtained for the short 15-residue dimers) reflect the fact that the local CC geometry is quite conserved across all experimental structures. A reconstruction with the top three PCs results in a median RMSD of 0.39Å, and a reconstruction with the full extended set of parameters (eq. 2) results in a median RMSD of 0.21Å.

Importantly, the structural variations introduced by the top three PCs are readily interpretable. The first PC corresponds to the axial rotation of an α-helix which also changes the Crick angles of all residues with respect to the CC axis (Fig. 3d). The second PC corresponds to an axial shift of the two α -helices relative to each other (Fig. 3e). The third component contains a variation of both CC radius and pitch, whereby an increase in pitch correlates with an increase in radius (Fig. 3f). The fourth and following components represent more complex movements.

## 6. Performance of PC-based parameterization

Finally, we have addressed the efficiency of the proposed CC parameterization. This analysis used a non-redundant collection of 151 canonical CC dimers with 23 residues per chain or longer extracted from the CC+ database [10] that had previously been used for a similar validation in [5]. Each structure from this collection has been optimally parameterized using the following procedure.

First, we align the average 15-residue fragment of the *a*-cluster described above with the target experimental structure in all possible registers (Fig. 4a). Expectedly, for a structure with a regular heptad repeat low RMSDs are obtained for every seventh alignment. Partial alignment to terminal fragments is provisioned for the cases when the structure starts or ends with positions other than *a*. This way a set of optimal placements of the average 15-residue dimer on the target structure is produced. Thereafter, for each such placement, the best fit upon parameterization with *k* top PCs (Fig. 3a) is obtained, and the mean values of the optimized parameters across all placed fragments are calculated. Finally, each placed fragment is transformed using the same set of parameters, and the resulting fragments are merged together along a straight line corresponding to the CC axis. At the end the backbone RMSD100 [15], rather than an ordinary RMSD, between the final model and the target structure is calculated to enable a direct comparison with related methods.

**Figure 4:**
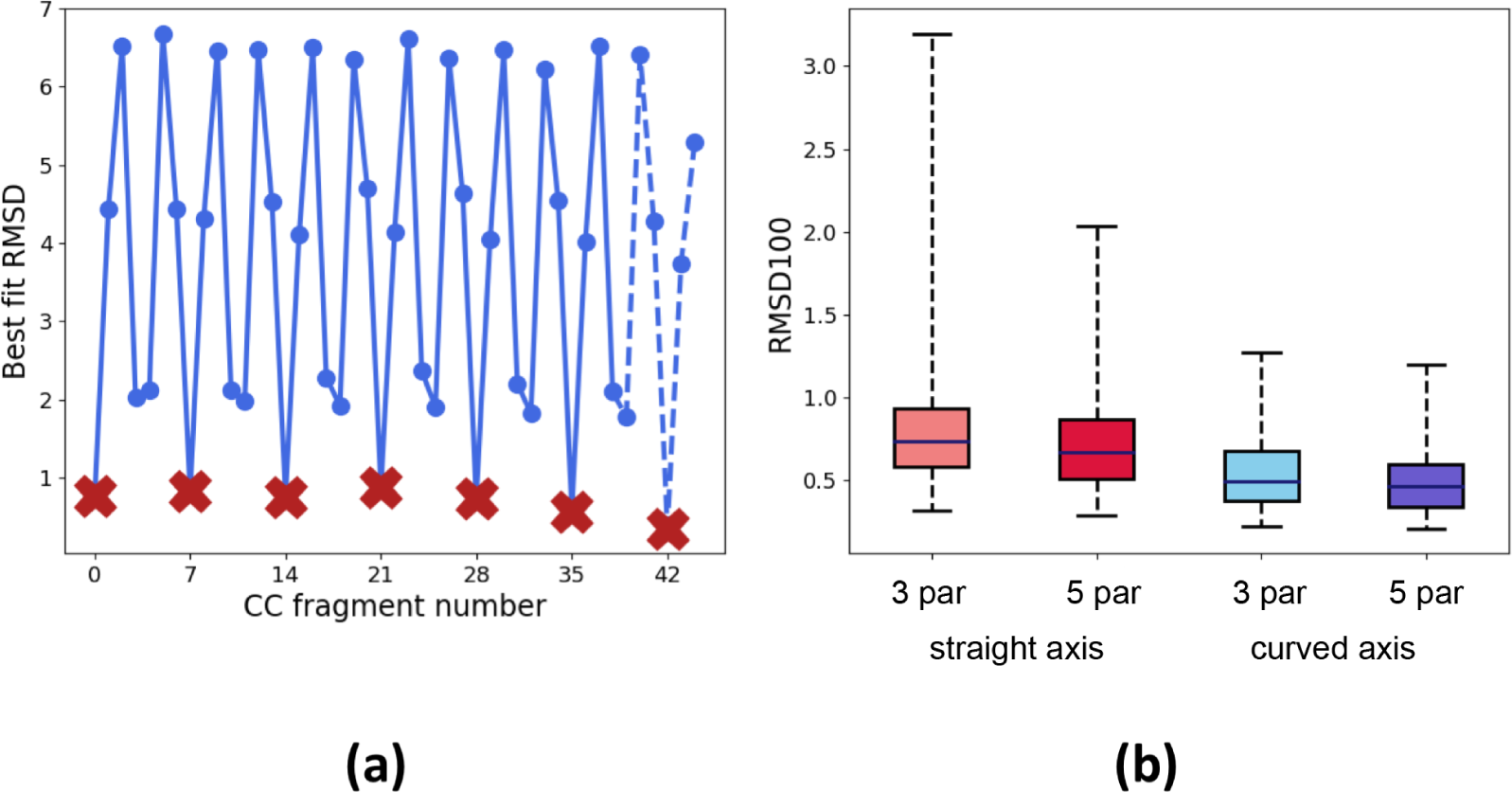
Parameterization of longer CCs. (a) Result of a step-wise alignment of 15-residue CC fragments from the a-cluster with a 44-residue experimental structure. Partial alignments are plotted with dashed lines. Selected fragments, indicated with red crosses, are fitted with a single parameter set and merged. (b) Box plots of the RMSD100 distributions for the validation set, fitted with a varying number of parameters.

Reconstruction of the validation set (Fig. 4b) using only three parameters and a straight CC axis yields a median RMSD100 of 0.74Å, while the maximal RMSD is 3.2Å. Not surprisingly, the largest deviations are observed for longer experimental structures which often have a pronounced bending of the CC axis. Such bending is often observed in heterodimeric CCs or in structures where the CC is interacting with another domain. It can also be due to lattice contacts within the crystal. Solely for the purposes of evaluating our parameterization algorithm, we have additionally introduced bending of the reconstructed CC to follow the CC axis of the experimental structure. In this case the median RMSD100 of the 3-parameter reconstruction decreases to 0.5Å (Fig. 4b).

## 7. Implementation

The program is implemented as a Python script. Interactive visualization is possible in PyMOL [16] with the help of the PyMOL-PyRosetta server link [17]. Protein Data Bank (PDB) files are processed with Biopython [18]. PyRMSD [12] is used for structural superpositions and multiple alignment. A standard minimizer of SciPy [19] was used to fit the parameterized models to the target structures.

The source code of the program can be freely downloaded at https://github.com/biocryst/CCParams.

## 8. Discussion

Modelling CC structures with a small number of parameters is important towards applications that require efficient sampling of all plausible CC geometries. One example is macromolecular X-ray crystallography, where experimental phases of reflections are often not available. In this case, one can estimate the phase using an appropriately placed initial structural model (this procedure is called ’molecular replacement’ (MR), [20]). A caveat here is that the MR procedure for CCs, being both elongated and repetitive structures, can be very challenging [21]. This demands a very accurate search model. Here, parameterization provides a means to efficiently sample all possible CC conformations. For instance, parameterization of CCs can be incorporated in a so-called ’evolutionary’ MR protocol [22]. Here, sampling of the rotational and translational space (six dimensions) could be done simultaneously with sampling just two or three parameters describing the CC geometry.

To this end, it is desirable to use a minimal number of parameters that provide the required accuracy of modelling. Importantly, the procedures proposed here use PC analysis and as a result provide for a highly efficient CC parameterization. In particular, the obtained parameters are orthogonal and correctly ranked, meaning that a sufficiently accurate reconstruction can be achieved with a minimal number of parameters. Indeed, our three-parameter reconstruction of the test set of CC structures (Fig. 4b) yields a median RMSD100 of 0.5Å with respect to experimental coordinates. This is somewhat better than the reconstruction achieved for a similar set of structures by the ISAMBARD algorithm [5]. The latter uses CC radius, CC pitch and the Crick angle as the three reconstruction parameters. The three top-scoring parameters obtained by our procedure (Fig. 4d-f) exhibit a correlated variation of the axial shift in addition to the same three classical Crick parameters. Being data-driven they provide a more efficient sampling of possible geometries.

Here it should be noted that the performance of the proposed parameterization critically depends on the starting collection of experimental structures. An efficient parameterization can be achieved only if such a collection samples the continuum of possible geometries well. For dimeric and trimeric CCs, this is likely to be the case. For CC that are not significantly represented in experimental databases, the derived parameterization may be less useful. In these cases unconstrained parametric models, or even general-purpose protein structure prediction methods are preferable.

In general, modelling a CC using a single ‘global’ set of parameters is advisable only for structures where a more or less regular geometry can be expected. This typically implies the presence of a regular sequence pattern such as the heptads. At the same time, longer CCs, even those based on regular patterns, will naturally have some variations especially in their CC pitch value that accumulate along their length. As the result, a reconstruction of a longer CC using a global set of parameters is less likely to achieve a sufficiently low RMSD with respect to the target structure, compared to shorter CCs. Furthermore, longer experimental CCs often reveal deviations of the CC axis from a straight line. Our analysis suggests that such deviations often represent the main source of discrepancy between the reconstruction and the real structure.

At the same time, many CC structures include discontinuities in the sequence pattern [23] which result in a pronounced local distortion of the CC geometry. Not unexpectedly, such structures are poorly suited for *de novo* modelling using a global set of parameters. In such cases, more sophisticated approaches should be used that take such irregularities into account. The available tools include a specialized fold-and-dock protocol of the popular Rosetta suite [24] and our CCFold program [11], which implement data-driven modelling algorithms designed for CC structures in particular.

